# USP7 inhibition perturbs proteostasis and tumorigenesis in triple negative breast cancer

**DOI:** 10.1101/2025.01.28.635372

**Authors:** Ahhyun Kim, Priya Gopalakrishnan, Claire C. Chen, Nikita Umesh, Angie Mordant, Natalie K. Barker, Laura E. Herring, Marina Suárez-Pizarro, Rasha T. Kakati, Philip M. Spanheimer, Michael J. Emanuele, Claudia A. Benavente

## Abstract

The deubiquitinase USP7 is a critical regulator of tumorigenesis, known for stabilizing the MDM2-p53 pathway. Emerging evidence highlights USP7’s p53-independent roles in proliferation and tumorigenesis. Triple negative breast cancers frequently inactivate p53 and this disease subtype remains difficult to treat and in need of new therapeutic options. Our study reveals that USP7 is upregulated in TNBC patient tumors. Importantly, genetic and pharmacologic USP7 inactivation impaired tumor progression in TNBC models. To explore USP7’s role in p53-mutant TNBCs, we performed deep quantitative proteomics across TNBC cell lines, identifying shared USP7 targets involved in cell proliferation, genome stability, and proteostasis. Acute USP7 inactivation allowed us to infer proximally controlled proteins which are likely direct targets. Surprisingly, many of the proteins downregulated by USP7 inhibition are E3 ubiquitin ligases. Thus, a key USP7 function in TNBC is to antagonize the degradation of ubiquitinating enzymes, since these enzymes are often susceptible to auto-ubiquitination and degradation. Notably, we identified TOPORS, a dual ubiquitin- and SUMO-ligase, among novel USP7 substrates. TOPORS interacts with the BRCA1-A DNA damage repair complex suggesting a USP7-TOPORS-BRAC1-A axis that might further explain the continued proliferation of genomically unstable TNBCs. Collectively, these data nominate USP7 as a potential therapeutic in TNBC.

## INTRODUCTION

Triple-negative breast cancer (TNBC) is characterized by the absence of estrogen and progesterone receptors (ER and PR) and the lack of amplification of Her2/ERBB2/Neu receptor tyrosine kinase. TNBCs largely overlap with the basal intrinsic molecular breast cancer (BBC) subtype (TCGA 2012). TNBC/basal subtype is an independent predictor of mortality, and this association has been refractory to advances in breast cancer therapy(Sørlie et al. 2001; Bhardwaj et al. 2023). TNBC tumors lack specific mutant oncoproteins (e.g., RAS, PI3K) but exhibit universal loss of the *TP53* tumor suppressor, leading to widespread genomic rearrangements (Spellman et al. 2012). Further, TNBCs exhibit functional loss of the retinoblastoma tumor suppressor (*RB1*), rendering CDK4/6 inhibitors also ineffective for TNBC treatment (Spellman et al. 2012; Finn et al. 2009). PARP inhibitors have been shown to be effective for TNBC patients with a BRCA1 mutation (5-10% of cases), and more recently the addition of immune checkpoint inhibitors in addition to chemotherapy have shown efficacy for metastatic TNBC and in the neoadjuvant setting for some patients with early TNBCs (Tung and Garber 2022; Cortes et al. 2022; Schmid et al. 2024). There is an unmet clinical need for better therapeutic options.

Ubiquitination is a three-step enzymatic cascade wherein E3 ubiquitin ligases modify substrates with the small protein ubiquitin. Polyubiquitination often leads to substrate degradation, making ubiquitin a central regulator in various physiological processes and diseases (Varshavsky 2012). Ubiquitination is a key regulator of various cellular functions, including gene expression and cell cycle progression. Ubiquitin can be removed from substrates by catalytic proteases termed deubiquitinases (DUBs) (Komander et al. 2009). Since DUBs remove ubiquitin from substrates, this prevents their degradation by the proteasome, often increasing the abundance of their targets post-transcriptionally. Among the 100 identified DUBs, ubiquitin-specific protease 7 (USP7) is the most extensively studied (Schauer et al. 2020). USP7 is essential for stabilizing proteins involved in critical cellular activities, including DNA replication, cell cycle regulation, and apoptosis. In many cancers, USP7 is overexpressed, stabilizing proteins crucial for tumor survival (Saha et al. 2023). Most research on USP7 has centered on its role in regulating TP53 and its E3 ligase, MDM2 (Chauhan et al. 2012; CHENG et al. 2013; Fan et al. 2013; Zhao et al. 2014), though in at least 50% of tumors, *TP53* is inactivated by mutations (Chen et al. 2022). Recent studies suggest USP7 may be a target of interest in cancers harboring *TP53* loss-of-function mutations, including triple negative breast cancer (TNBC), where FOXM1 was identified as TP53*-*independent target of USP7 (Yi et al. 2023).

In this study, we investigated the role of USP7 in TNBC progression, focusing on its impact on tumor growth, metastasis, and downstream regulatory networks. Using genetic and pharmacological approaches, we explored how USP7 inhibition affects TNBC tumors both in vitro and in vivo. Through integrative proteomic and transcriptomic analyses, we identified key pathways and substrates regulated by USP7, including proteins involved in ubiquitination and DNA repair. Our findings reveal novel insights into USP7’s contributions to TNBC pathogenesis and highlight its potential as a therapeutic target for this aggressive breast cancer subtype.

## RESULTS

### Recurrent USP7 amplification and overexpression is linked to poor prognosis in breast cancer

Elevated levels of USP7 have been observed in many cancers. Analysis of breast cancer patient data from cBioPortal and TCGA indicate that *USP7* is recurrently amplified at the chromosomal level and significantly overexpressed (Figure 1A-B). USP7 is among the most recurrently upregulated DUBs in breast cancer with nearly a quarter (24%) of patient tumors showing genomic amplification or mRNA overexpression of *USP7*, a rate higher than the well-known oncogene *MYC* (22%), as well as other cancer associated DUBs, including USP22, USP2, USP28 and BAP1 (Figure 1A). Interestingly, USP7 overexpression occurs across different breast cancer subtypes, except Claudin-low tumors (Supplemental Figure 1). We also analyzed breast cancer cells sensitivity to USP7 KO using the large-scale functional genomic screens available on the Cancer DepMap after sub-classifying them into known pathologic subtypes (ER+/Her2-, Her2+, and TNBC; Figure 1C). This classification scheme is highly predictive of sensitivity to cell cycle inhibitors targeting CDK4/6, with TNBC cells being universally resistant (Finn et al. 2009). TNBC cell lines are sensitive to USP7 KO, despite their recurrent loss of p53 (Figure 1C). This supports the notion that USP7 could contribute to tumorigenesis through mechanisms which are independent of p53.

**Figure 1.**
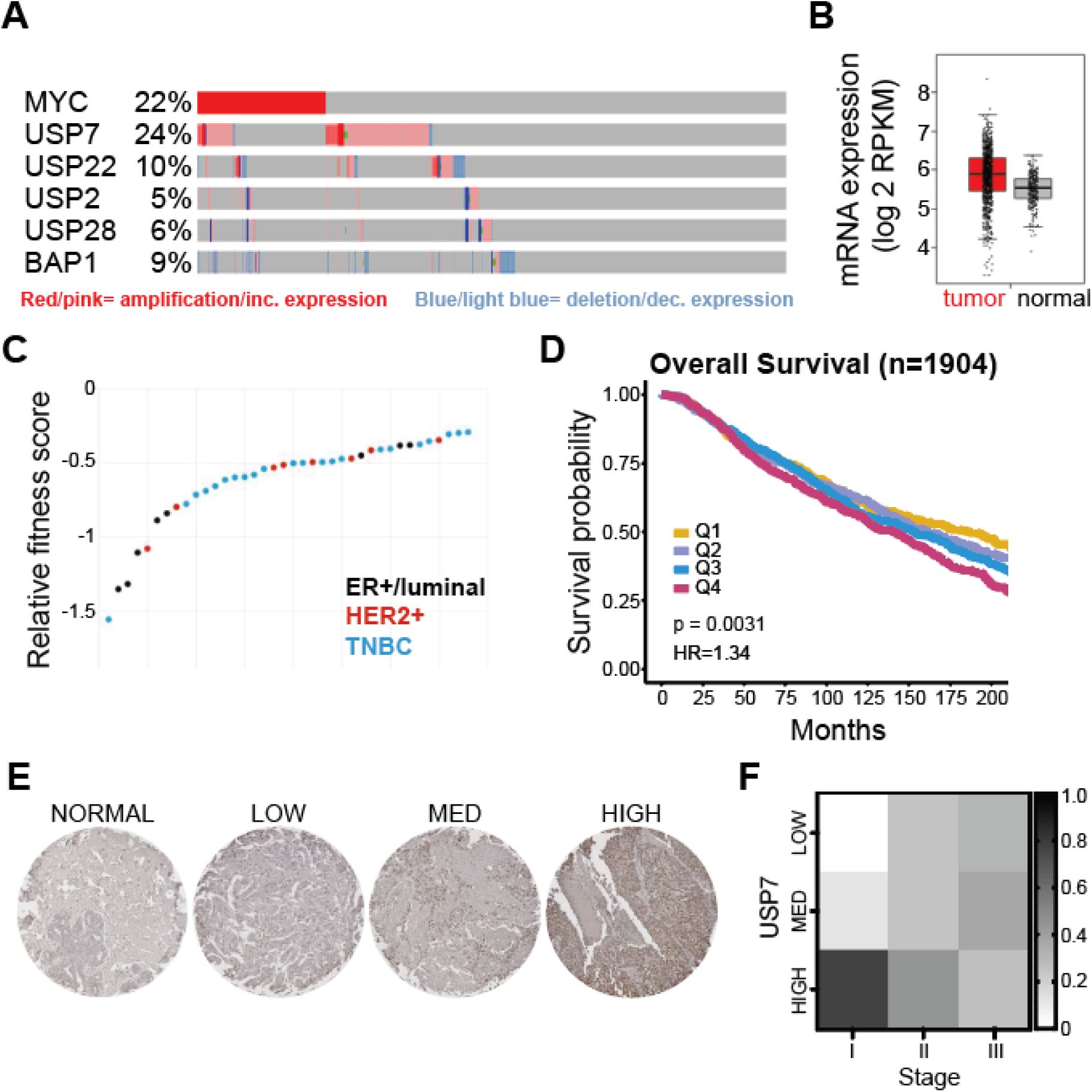
USP7 overexpression is associated with poor prognosis in breast cancer. (A) cBioportal data for breast cancer tumors shows USP7 is recurrently amplified and overexpressed in breast cancer. Red=amplification; pink=increased expression; blue=deletion; light blue=decreased expression. (B) TCGA breast cancer database shows USP7 mRNA expression is elevated across all breast cancer tumors (n=1085, red) compared to normal breast tissue (n=291, grey). (C) Analysis of DepMap data on the effect of USP7 CRISPR KO on breast cancer cell fitness/proliferation separated by pathologic breast cancer subtypes. (D) Overall survival analysis from the METABRIC breast cancer database shows high USP7 expression correlates with poorer survival (n=1904). Patients grouped into quartiles based on USP7 expression: Q1 (yellow)=lowest USP7 expression; Q4 (magenta)=highest USP7 expression. Log-rank test p=0.0031, HR=1.34. (E) Representative IHC images of core human tissues from a TNBC tissue microarray stained for USP7. (F) Heat map showing percentage of core tissues with low, medium, or high USP7 protein levels, subdivided into TNBC stage I, II, or III.

Consistent with a potential role in disease, METABRIC analysis of 1904 breast cancer patients revealed that those with the highest levels of USP7 (top quartile, Q4) had the worst overall survival compared to patients with lower levels of USP7, with Q1 representing the lowest levels (Figure 1D).

Since proteins in the ubiquitin pathway are often regulated post-translationally, we examined USP7 protein levels. Significantly, our analysis of USP7 protein expression in TNBC using a patient tissue microarray with 111 TNBC tumors, found that 55% of TNBC tumors had medium-to-high USP7 protein expression. Elevated USP7 protein levels were observed across all stages of this invasive breast cancer (Figure 1E-F). These findings indicate that USP7 is nearly universally overexpressed in TNBC and negatively associates with patient outcomes.

### USP7 promotes TNBC clonogenicity

To investigate how elevated USP7 in TNBC increases malignancy, we selected three TNBC cell lines – SUM159, MDA-MB-231, and MDA-MB-468 – all of which were confirmed to express varying degrees of USP7 compared to the immortalized normal breast cell line MCF10A. We included the non-TNBC breast cancer cell line, MCF7, as a positive control (Figure 2A) (Schauer et al. 2020). Using CRISPR/Cas9, we generated syngeneic *USP7* knockout (KO) clones and non-targeting control clones for each of the three TNBC cell lines (Figure 2B and Supplemental Figure 2). Proliferation analyses revealed a moderate but consistent increase in doubling time in USP7 KO clones compared to the control in the three TNBC lines (Supplemental Figure 2). Of significance, USP7 KO conferred a significantly reduced ability to form colonies in clonogenic assays, with an average reduction of 94.0 ± 10.3% in the number of colonies formed compared to the control across the three cell lines (Figure 2C). MDA-MB-231 and MDA-MB-468 cells lacking USP7 were almost completely unable to form colonies.

**Figure 2.**
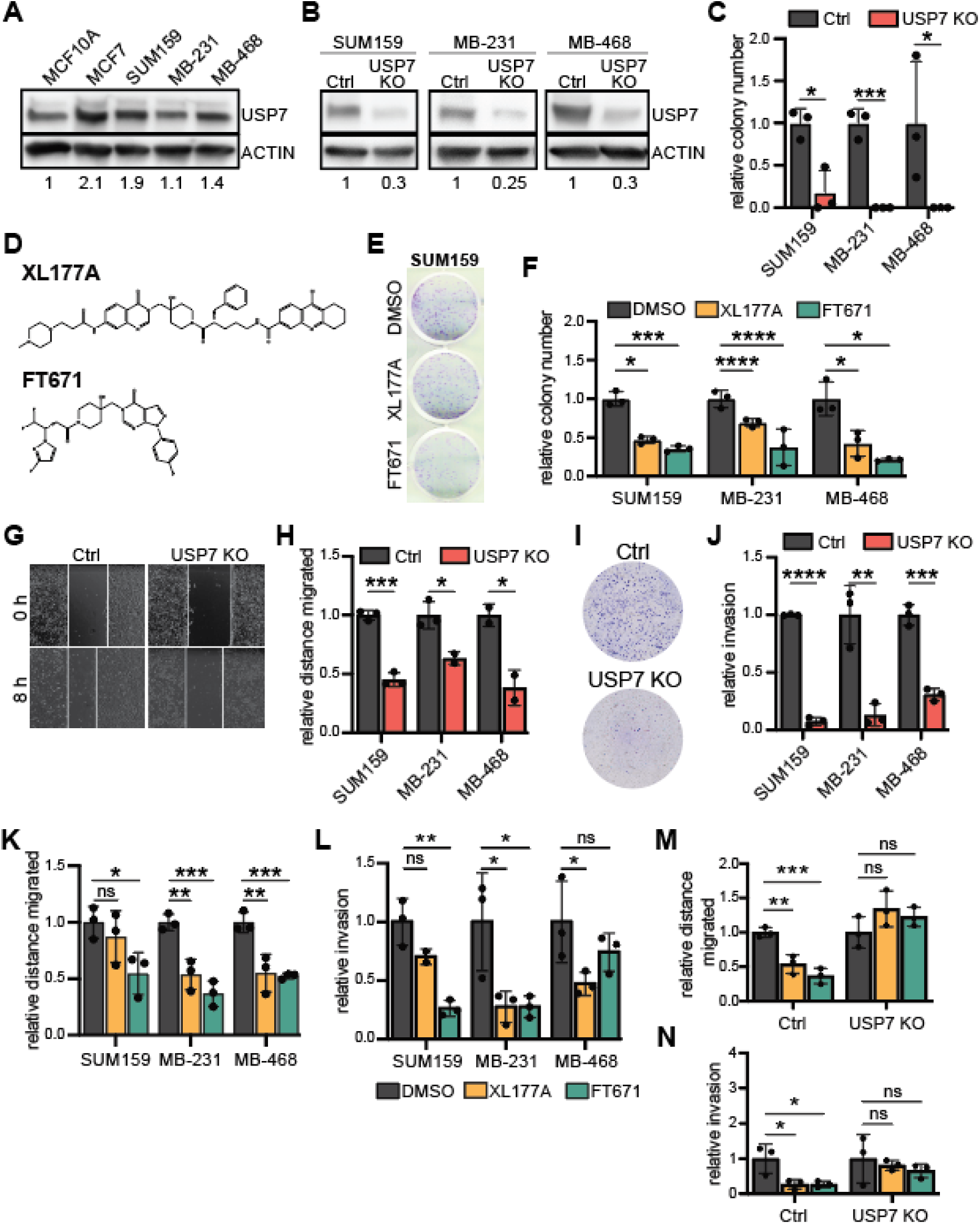
USP7 promotes TNBC clonogenicity and cell migration and invasion. (A) Western Blot analysis showing USP7 protein expression levels in malignant breast cancer cells, including TNBC, compared to the non-malignant MCF10A mammary epithelial cell line used as control. Quantification relative to MCF10A expression shown at the bottom after β-actin normalization. β-actin was used as loading control. (B) Representative Western blot analysis of USP7 in CRISPR/Cas9-mediated USP7 knockout (USP7 KO) clones of TNBC cell lines in comparison to non-targeting vector control (Ctrl). β-actin was used as loading control. (C) Bar graph of colony counts from clonogenic assays comparing USP7 KO to Ctrl. (D) Chemical structures of USP7 inhibitors that were used for in vitro assays. (E) Representative images of colonies stained with crystal violet comparing SUM159 treated with XL177A or FT671. DMSO used as vehicle control. (F) Bar graph of colony counts from clonogenic assays comparing TNBC cells treated with XL177A or FT671. DMSO as control. (G) Representative image from scratch-wound assay comparing wound closure of SUM159 non-targeting vector control (Ctrl) and USP7 KO clone over 8 hours. White lines represent the wound edge. (H) Quantification of the distance cells migrated in scratch-wound assays for each TNBC cell line. (I) Representative images from Transwell invasion assays with cells stained with crystal violet comparing SUM159 Ctrl and USP7 KO clone. (J) Quantification of the number of cells invaded across a Transwell membrane for each of the TNBC cell lines. (K) Quantification of the distance cells migrated for TNBC cells treated with XL177A or FT671. DMSO used as vehicle control. (L) Quantification of cell invasion for TNBC cells treated with XL177A or FT671. DMSO used as vehicle control. (M) Quantification of the distance cells migrated normalized to DMSO for MDA-MB-231 Ctrl and USP7 KO. (N) Quantification of cell invasion normalized to DMSO for MDA-MB-231 Ctrl and USP7 KO. For all bar graphs, values were normalized to control. Each bar is shown as mean ± SD, with each dot representing an independent replicate, n= 3. For all graphs: ns=not significant, *p<0.05, **p<0.01, ***p<0.001, ****p<0.0001 by unpaired two-tailed t test.

Highly selective USP7 small molecule inhibitors have been extensively developed with nanomolar potency. For in vitro USP7 inhibition analyses, we used XL177A, an irreversible inhibitor with an IC50 of 0.34 nM (Schauer et al. 2020), and FT671, a noncovalent inhibitor with an IC50 of 52 nM (Turnbull et al. 2017) (Figure 2D). Both inhibitors have been screened against a panel of other DUBs and both showed high specificity for USP7 inhibition (Schauer et al. 2020; Turnbull et al. 2017). USP7 pharmacological inhibition had minimal effects on cell viability and cell proliferation in vitro (Supplemental Figure 3). However, similar to USP7 KO, treatment with XL177A or FT671 led to a significant reduction of 47.1 ± 14.4% and 68.5 ± 8.7%, respectively, in colony numbers across the three cell lines compared to DMSO (Figure 2E-F).

These findings suggest that while USP7 has only a modest impact on cell proliferation in TNBC cells lines, it plays a critical role in promoting cell stemness, essential for long-term tumor growth and survival.

### USP7 promotes TNBC cell migration and invasion

Metastasis is the primary cause of breast cancer-related mortality and recurrence. USP7 plays a critical role in stabilizing proteins involved in metastasis (Duan et al. 2022; Liu et al. 2020). A recent study also reported that USP7 promotes breast cancer migration and invasion, including in TNBC, through stabilization of the epigenetic reader ZMYND8 (Tang et al. 2024). In line with these observations, USP7 KO resulted in a significant decrease in cell migration in scratch-wound healing assays, with an average reduction of 47.2 ± 19.5% across the three TNBC cell lines (Figure 2G-H). Invasion assays using Matrigel-coated inserts also revealed a significant decrease in invasiveness of 82.8 ± 12.1% in USP7 KO cells compared to control across all three TNBC cell lines (Figure 2I-J).

Pharmacological inhibition of USP7 with FT671 similarly lead to reduced cell migration in all three cell lines, while XL177A slowed migration in two out of three (Figure 2K). USP7 inhibition also reduced the number of invaded cells compared to DMSO-treated controls in MDA-MB-231 (Figure 2L). Although the reduction in invasion with USP7 inhibitors was less pronounced for SUM159 and MDA-MB-468, we observed a trend toward decreased invasion with USP7 inhibition (Figure 2L). Confirming the on-target effects of both inhibitors, USP7 KO cells treated with USP7 inhibitors resulted in minimal changes in cell migration and invasion compared to control (Figure 2M-N and Supplemental Figure 3).

### USP7 drives tumor growth and metastasis in TNBC

Building on the observed effects of USP7 on cell stemness, migration, and invasion in TNBC, we next evaluated its role in promoting in vivo TNBC tumor growth and metastasis. NSG mice were orthotopically injected with MDA-MB-231, either control or USP7 KO, both expressing luciferase for bioluminescent monitoring. Tumors were tracked biweekly for 10 weeks. Tumor growth in the USP7 KO group was significantly slower than the control group. At the study endpoint, the USP7 KO group showed a significantly reduced tumor burden, based on bioluminescent imaging, and no detectable metastasis, whereas the control mice exhibited larger and highly metastatic tumors (Figure 3A). Delayed tumor growth was also observed in MDA-MB-468-derived USP7 KO tumors (Supplemental Figure 4A).

**Figure 3.**
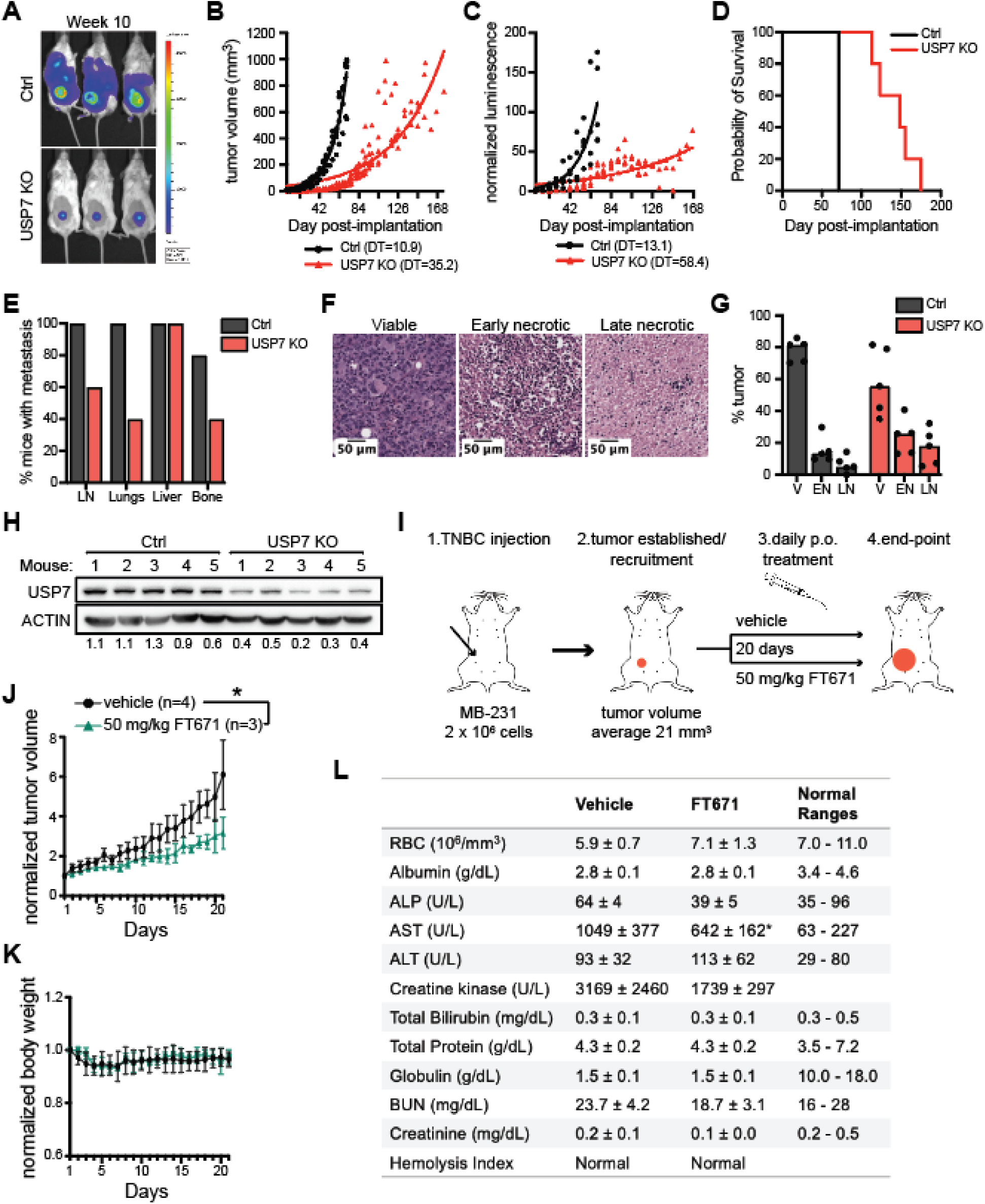
USP7 promotes TNBC tumor growth and metastasis. (A) Representative bioluminescent images of non-targeting vector control (Ctrl) and USP7 KO primary tumors in the mice at week 10. Overall intensity of bioluminescence shown by area and color, with red having the highest photon count and blue the lowest. (B) Tumor growth curve comparing MDA-MB-231 Ctrl (n=5; black) and USP7 KO (n=5; red). Groups reached endpoint when average tumor volume was 1000 mm^3^ measured by caliper. DT=doubling time in days. (C) Tumor growth curve comparing MDA-MB-231 Ctrl (black) and USP7 KO (red) measured by bioluminescence. DT=doubling time in days. (D) Survival curve comparing Ctrl and USP7 KO. Mice had reached endpoint based on final average tumor volume of 1000 mm^3^. (E) Quantification of percentage of mice bearing metastasis at endpoint. Black=Ctrl (n=5); Red=USP7 KO (n=5). LN=lymph nodes. (F) Representative histology sections of hematoxylin and eosin stained tumors distinguishing viable, early necrotic, and late necrotic tissue areas. (G) Percentage of viable, early necrotic, or late necrotic tumor areas comparing Ctrl to USP7 KO. (H) Western blot analysis for USP7 protein levels in Ctrl versus USP7 KO primary tumors. USP7 values normalized to control average after β-actin normalization. (I) Schematic workflow of mice bearing MDA-MB-231 tumors administered USP7 inhibitor, FT671, by oral gavage daily for 20 days. (J) Normalized tumor growth comparing vehicle (n=4; black) and FT671 (n=3; green) over 20 days of treatment. *p < 0.05 by two-way ANOVA. (K) Daily average mouse body weight over 20 days of treatment for vehicle (n=4; black) and FT671 treated (n=3; green) mice. (L) Blood chemistry in the vehicle mice (*n*=4) versus mice treated with 50 mg/kg FT671 (n=3). Blood was collected after 20 days of daily treatment.

To account for differing tumor growth rates in assessing metastasis, a follow-up study set the endpoint to when tumor volumes reached 1000 mm^3^. USP7 KO tumors grew significantly slower, taking over 25 weeks to reach the endpoint compared to 10 weeks for control tumors (Figure 3B-C). This delay in tumor growth translated to a significantly extended survival period for the USP7 KO group (Figure 3D). Importantly, even when tumors reached comparable sizes, mice in the USP7 KO group exhibited significantly fewer metastatic lesions in lymph nodes, lungs, and bones compared to the control group (Figure 3E). Interestingly, liver metastases were observed in all mice at comparable levels, suggesting that liver metastasis may be driven by USP7-independent factors such as chemokines and inflammatory signals (Ma et al. 2015).

Transcriptomic analysis using bulk RNA-seq of MDA-MB-231 tumors revealed 1308 up-regulated and 1053 down-regulated genes in USP7 KO mice compared to controls (Supplemental Figure 4B). Gene ontology (GO) analysis showed that the most enriched biological process for upregulated genes was negative chemotaxis, consistent with our observed reduced migration, invasion, and metastasis (Supplemental Figure 4C). Downregulated genes, including TNF, were significantly associated with the negative regulation of extrinsic apoptotic signaling pathway in absence of ligand (Supplemental Figure 4D). Histological analysis further supported these findings, with hematoxylin and eosin staining showing increased necrosis in USP7 KO tumors (Figure 3F). These tumors displayed a significantly higher percentage of both early and late necrotic regions compared to controls (Figure 3G). Western blot analysis confirmed reduced USP7 protein levels in the USP7 KO tumors (Figure 3H). Altogether, these findings highlight USP7 as a critical driver of TNBC tumor growth and metastasis.

### Pharmacological inhibition of USP7 slows TNBC tumor growth

Our findings demonstrated that genetic targeting of USP7 significantly reduces tumor growth. Building on the specificity of USP7 inhibitors observed in vitro, we investigated the therapeutic potential of FT671 in vivo. To evaluate the effects of USP7 inhibition, luciferase-expressing MDA-MB-231 cells were orthotopically injected into the mammary fat pads of NSG mice. Once tumors were established, mice were randomly assigned to receive daily oral (p.o.) treatment with either vehicle or 50 mg/kg FT671 (Figure 3I). Tumor growth, measured daily by calipers, was significantly reduced in the FT671-treated group compared to the vehicle control over the 20-day treatment period (p=0.0312) (Figure 3J). Importantly, no differences in body weight were observed between the two groups, indicating that FT671 treatment was well tolerated (Figure 3K). Toxicological analyses of the liver, heart, and muscle revealed no additional toxicity from FT671 treatment beyond the effects associated with DMSO, known to induce liver fibrosis (Figure 3L). These results highlight the potential of USP7 inhibitors as a potentially safe and effective therapeutic strategy for slowing TNBC tumor growth.

### Proteomics analysis uncovers TOPORS as a novel USP7 substrate

To investigate the mechanisms underlying the poor outcomes associated with USP7 upregulation and the impact of USP7 inactivation of TNBC cells, we aimed to identify downstream ubiquitin signaling pathways leading to altered protein abundance that are affected by USP7. We employed TMT-based quantitative mass spectrometry to define the proteome-wide changes following USP7 inactivation. Three TNBC cell lines, MDA-MB-231, SUM159, and SUM149, were analyzed. All three were treated with either of the selective pharmacological USP7 inhibitors XL177A or FT671, or DMSO as a control, and profiled using multiplexed, quantitative proteomics using tandem mass tagging (Figure 4A) (Schauer et al. 2020; Turnbull et al. 2017). To reveal proximal changes resulting from the direct regulation of USP7, which are mostly likely to be USP7 substrates, rather than secondary downstream alterations (e.g., changes in transcription factor expression), an acute, 8-hour treatment duration was utilized for all cell lines and drug treatments. This approach allowed us to capture early proteomic alterations directly linked to USP7 activity.

**Figure 4.**
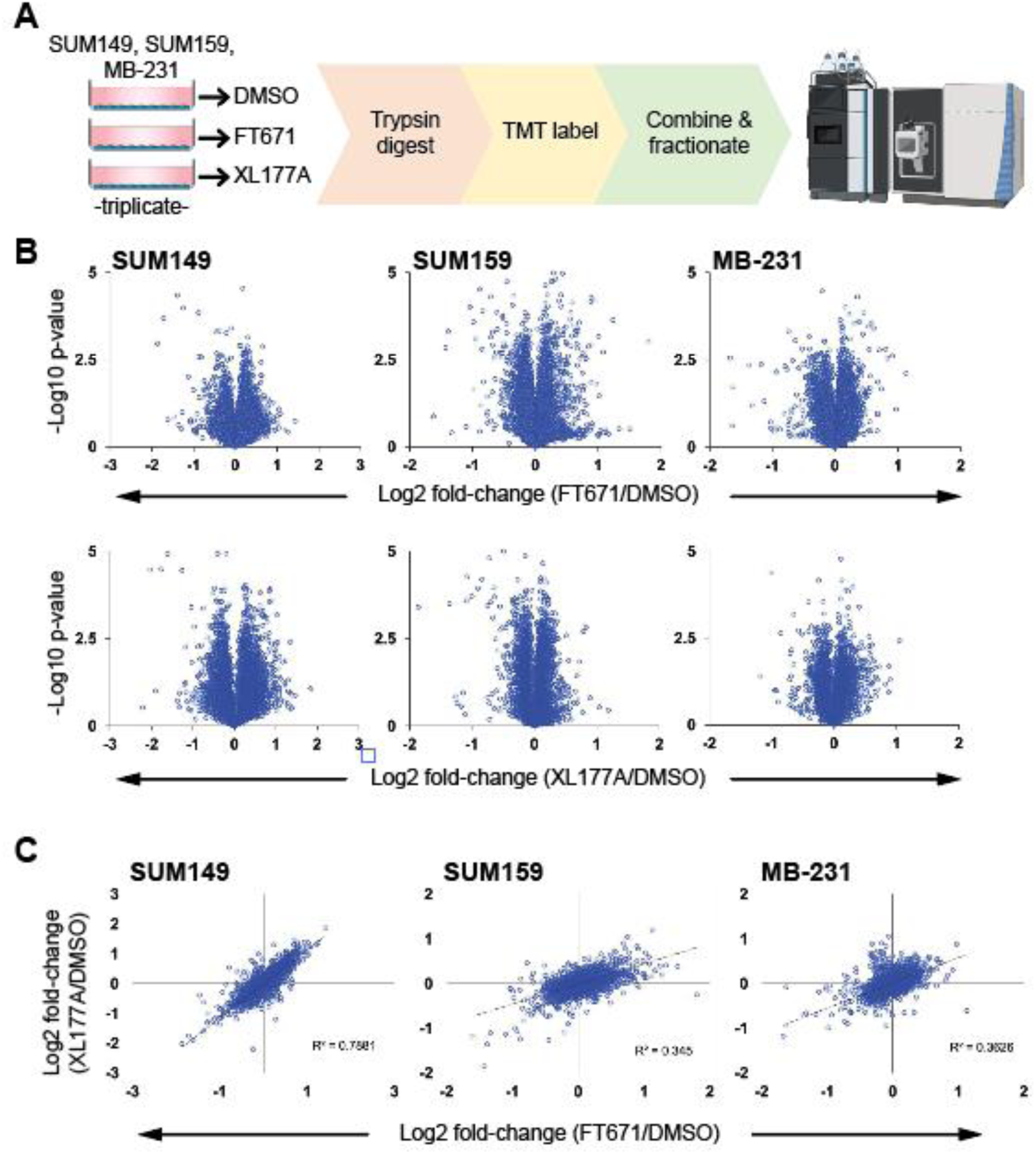
Deep, quantitative proteomics of USP7 inhibited TNBC cells. (A) Schematic describing the quantitative proteomics workflow for the USP7 inhibitor (USP7i) analysis. SUM149, SUM159, and MDA-MB-231 TNBC cell lines were grown and treated with DMSO, 1 μM FT671, or 1 μM XL177A for 8 h. Lysates are digested using trypsin and LysC peptides are TMT labeled, mixed, and fractionated. Peptides are analyzed by mass spectrometry (MS) and protein abundances determined. (B) Volcano plot showing the log2 fold-change (x axis) and -log10 p-value (y axis) of all proteins identified by MS when comparing samples from USP7i-treated (FT671 or XL1774A) cells to samples from DMSO-treated cells. Each dot represents an individual protein. (C) Plot showing the log2 fold-change ratio of all proteins identified by MS when comparing samples from FT671-treated to DMSO-treated cells (x axis) vs the log2 fold-change ratio of all proteins identified by MS when comparing samples from XL177A-treated to DMSO-treated cells (y axis).

We read deep into the proteomes of all three cell lines. After filtering for likely contaminants, we identified 7961 proteins in SUM149s, 8069 in SUM159, and 7845 proteins in MDA-MB-231. Collectively, we identified 8815 unique proteins species between the three cell lines. Among those, 7845 proteins (∼89%) were identified in more than one cell line and 7102 (80%) were identified in all three cell lines. The high concordance between cell lines underscores their similarity based on their shared disease subtype and tissue of origin.

All three cell lines treated were analyzed for quantitative changes in protein abundance using TMT labeling, comparing both FT671 and XL177A to DMSO control treated cells. Volcano plots for these 6 experimental conditions are shown in Figure 4B. Importantly, comparison of FT671 to XL177A treatment revealed a high concordance, underscoring the previously reported, on-target activity of both inhibitors towards USP7, particularly notable given that these two drugs lack structural similarity (Figure 4C).

Among the most significantly downregulated proteins were known USP7 substrates. This included UVSSA (Higa et al. 2016), FBXO38 (Georges et al. 2019), TRIP12 (Cai et al. 2015), and TRIM27 (Hao et al. 2015) (Figure 5A), although none have been previously reported in TNBC cell lines. We identified many other common hits that scored in multiple cell lines, many of which were E3 ligases. Among them was RNF220, a RING-finger ubiquitin ligase which we validated by western blot (Figure 5A and Supplemental Figure 5). L3MBTL2 is chromatin associated protein, and MGA is a dimerization partner with the oncogene MYC. Both were also consistently downregulated across cell lines treated with USP7 inhibitors (Figure 5A). Surprisingly, we identified several substrate receptor proteins for the CUL4-based RING E3 ubiquitin ligase (CRL4). This modular family of E3 ligases uses CUL4 as a backbone and the adaptor protein DDB1 to directly engage substrate receptors (see schematic in Figure 5B). The CRL4 substrate receptors are often referred to as DCAFS, for DDB1- and CUL4-associated factors. Among those downregulated in multiple cell lines were BRWD1/DCAF19, BRWD3, PHIP/DCAF14, and DCAF5 (Figure 5B). Collectively, the ability of USP7 to downregulate the abundance of many E3 ligases, including several DCAFs, following acute inactivation points to a broad role in coordinating proteostasis through the regulation of multiple E3 ligases.

**Figure 5.**
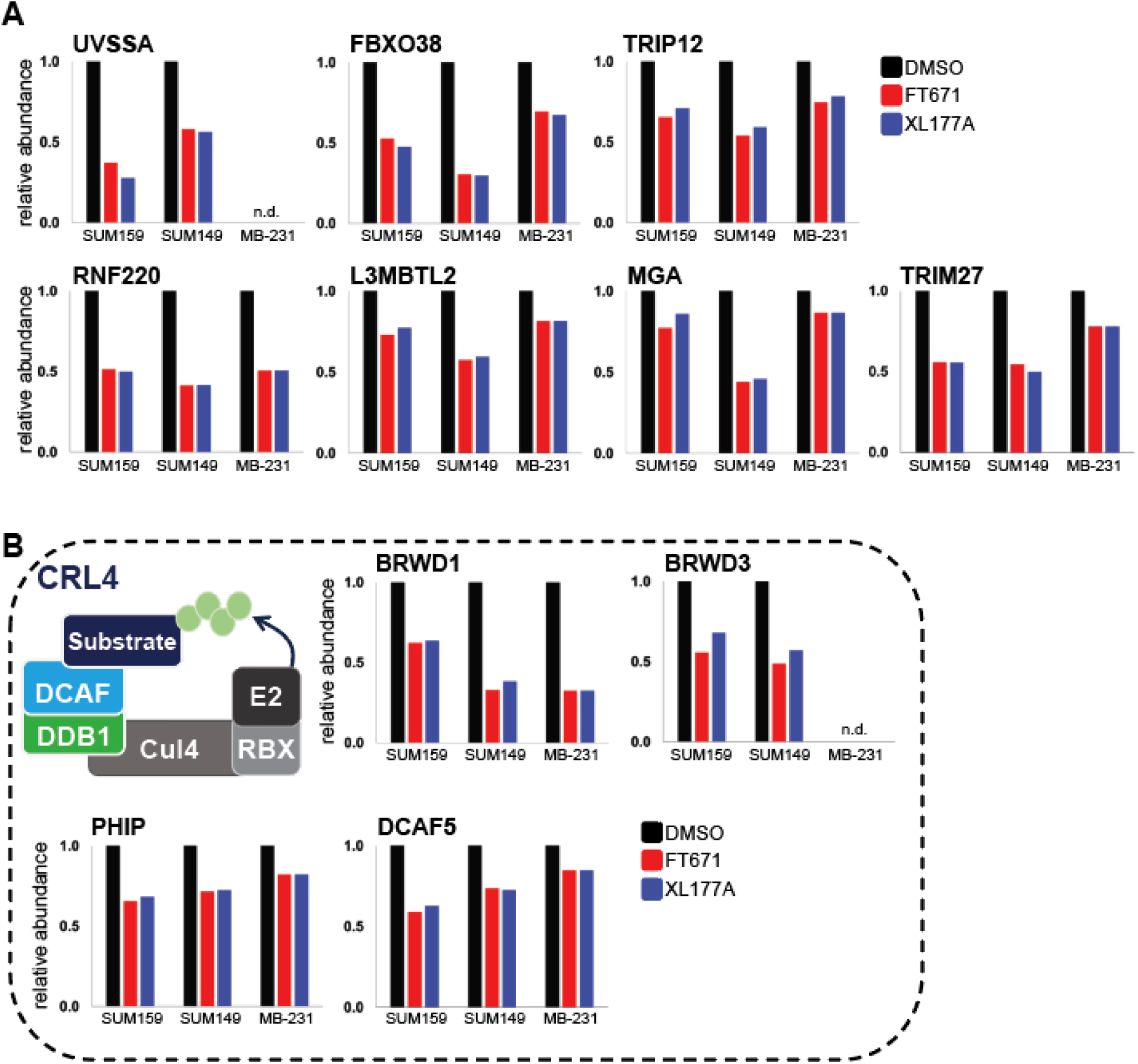
USP7 regulates diverse substrates, including several ubiquitin ligases. (A) Analysis of specific substrates from quantitative proteomics experiments preformed in three cell lines and with two distinct USP7 inhibitors. The known substrates UVSSA, FBXO38 and TRIP12 were all significantly downregulated in response to USP7 inhibition in multiple cell lines and using both inhibitors. IN addition, several other proteins were downregulated across multiple cell lines and including RNF220, L3MBTL2, MGA and TRIM27. (B) Several substrate receptors for the CRL4 ubiquitin ligase were similarly decreased by USP7 inhibition. A schematic of the modular CRL4 complex is shown. Interchangeable DDB1- and CUL4-associated factors (DCAFs) bridge substrates to the ubiquitin ligase complex by simultaneously binding DDB1 and ubiquitin targets. Among the substrate receptors controlled by acute USP7 inhibition are BRWD1, BRWD3, PHIP and DCAF5.

Among the most downregulated proteins in multiple cell lines, TOPORS is a multifunctional protein implicated in cell growth, proliferation, DNA damage response, and ubiquitination. It functions as an E3 ubiquitin ligase and potentially an E3 SUMO ligase (Denuc and Marfany 2010). While its role remains poorly characterized, recent studies suggest it is involved in relieving DNA-protein crosslinks (Carnie et al. 2024; Liu et al. 2024). However, its function in TNBC has not been explored. Not previously reported as a direct USP7 substrate, TOPORS was significantly downregulated by both USP7 inhibitors in both SUM159 and SUM149 cells (Figure 6A), although it was not detected in MDA-MB-231 cells. In SUM149 cells treated with FT671, TOPORS showed the most significant decrease among all 7961 proteins quantified and was the second most downregulated protein in SUM149 treated with XL177A (Figure 6B). We validated the downregulation of TOPORS by western blot in SUM159 cells and extended that analysis to show it was also downregulated in MDA-MB-231, where it was not detected by mass spectrometry (Figure 6C). The decrease in TOPORS was due to on target USP7 activity, as it was recapitulated by depletion of USP7 with multiple silencing RNAs (siRNA) (Figure 6D). In addition, this decrease was due to a destabilization of TOPORS, consistent with the notion that USP7 controls TOPORS ubiquitination and degradation, since USP7 inhibition with FT671 decreased its half-life in a cycloheximide chase assay (Figure 6E). We confirmed that this was direct by assaying the interaction between USP7 and TOPORS, which we could detect by reciprocal coIP in HEK293T cells (Figure 6F-G).

**Figure 6.**
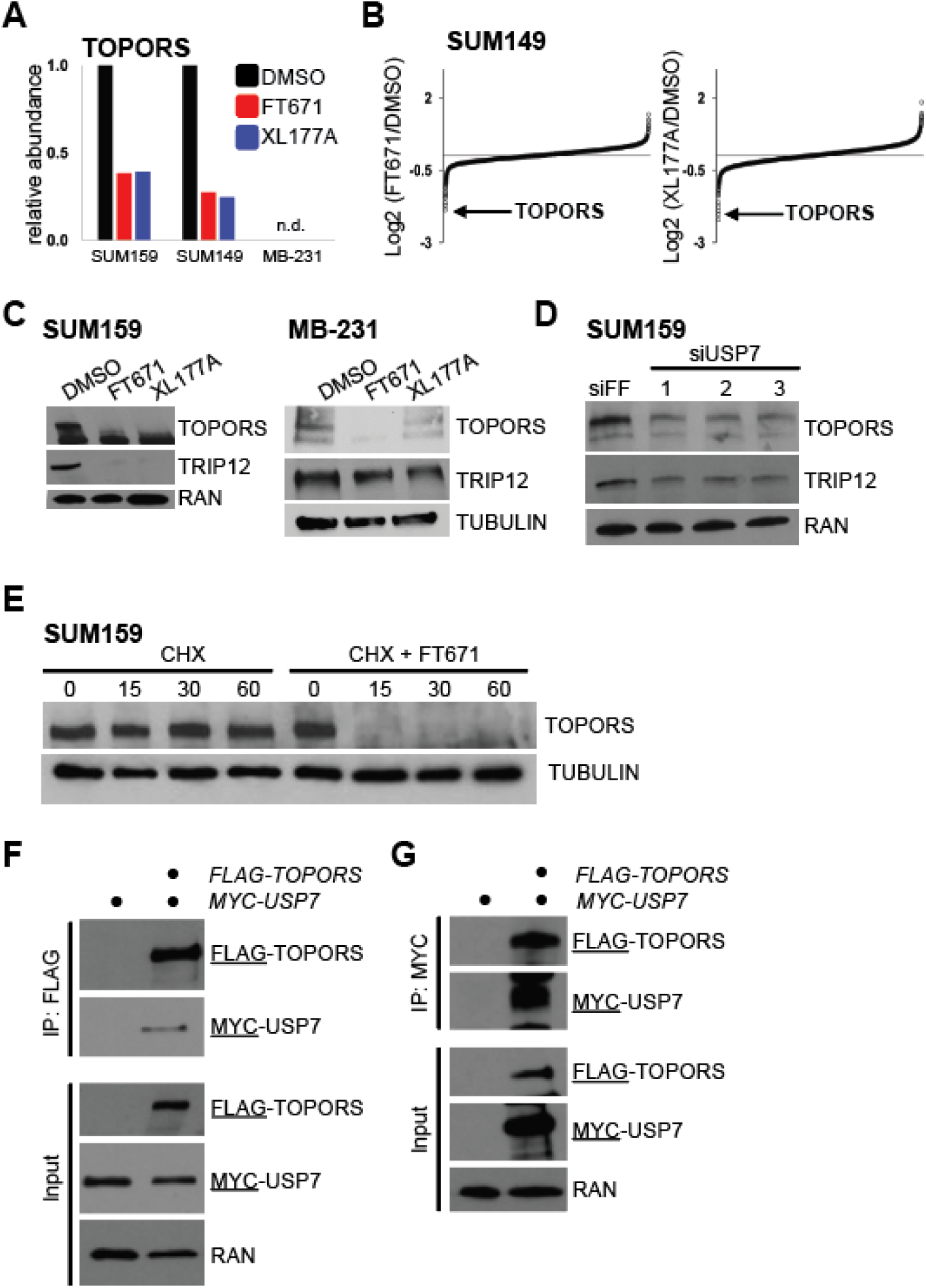
USP7 regulates TOPORS proteosomal degradation. (A) TOPORS was significantly downregulated by USP7 inhibition quantitative proteomic analysis in response to FT671 and XL177A treatment in SUM149 and SUM159 cell lines. (B) Plot showing the log2 ratio of all proteins identified by MS when comparing samples from FT671-(left) or XL177A-treated (right) to DMSO-treated SUM149 cells. Each dot along the x axis represent an individual protein. (C) SUM159 and MDA-MB-231 cells were treated with 1μM of either FT671 or XL177A and analyzed by western blot for TOPORS and the positive control TRIP12. (D) SUM159 cells were depleted of USP7 using siRNA and analyzed for the expression of TOPORS and TRIP12. (E) SUM159 cells were treated with cycloheximide to block translation, or cycloheximide the presence of FT671. The half-life of TOPORS was measured at the indicated time points, shown in minutes, following inhibitors addition. (F and G) HEK293T cells were transfected with indicated plasmids and analyzed by coIP for either FLAG-TOPORS (left) or MYC-USP7 (right).

### TOPORS associates with the BRCA1-A Complex

TOPORS has been linked to topoisomerase 1 (Rasheed et al. 2002) and has previously been shown to play a role in resolving DNA-protein crosslinks (Carnie et al. 2024; Liu et al. 2024). To gain deeper insight into its function, we carried out a protein interaction mass spectrometry screen (Figure 7A). FLAG-TOPORS was expressed in triplicate samples of HEK293T cells, and following lysis and immunoprecipitation, was analyzed by mass spectrometry. Control samples from cells expressing an empty vector were analyzed in parallel. USP7 was among the top scoring interactors, further validated their interaction and the direct regulation of TOPORS by USP7 (Figure 7B). Interestingly, several other DUBs were also identified, including USP11, USP30 and USP1. Several other interesting proteins were among the most enriched including SUMO1 and two SUMO deconjugating enzymes, SENP1 and SENP2. Surprisingly, topoisomerases 2A and 2B were both enriched, although we did detect an enrichment for topoisomerase 1 (Figure 7C).

**Figure 7.**
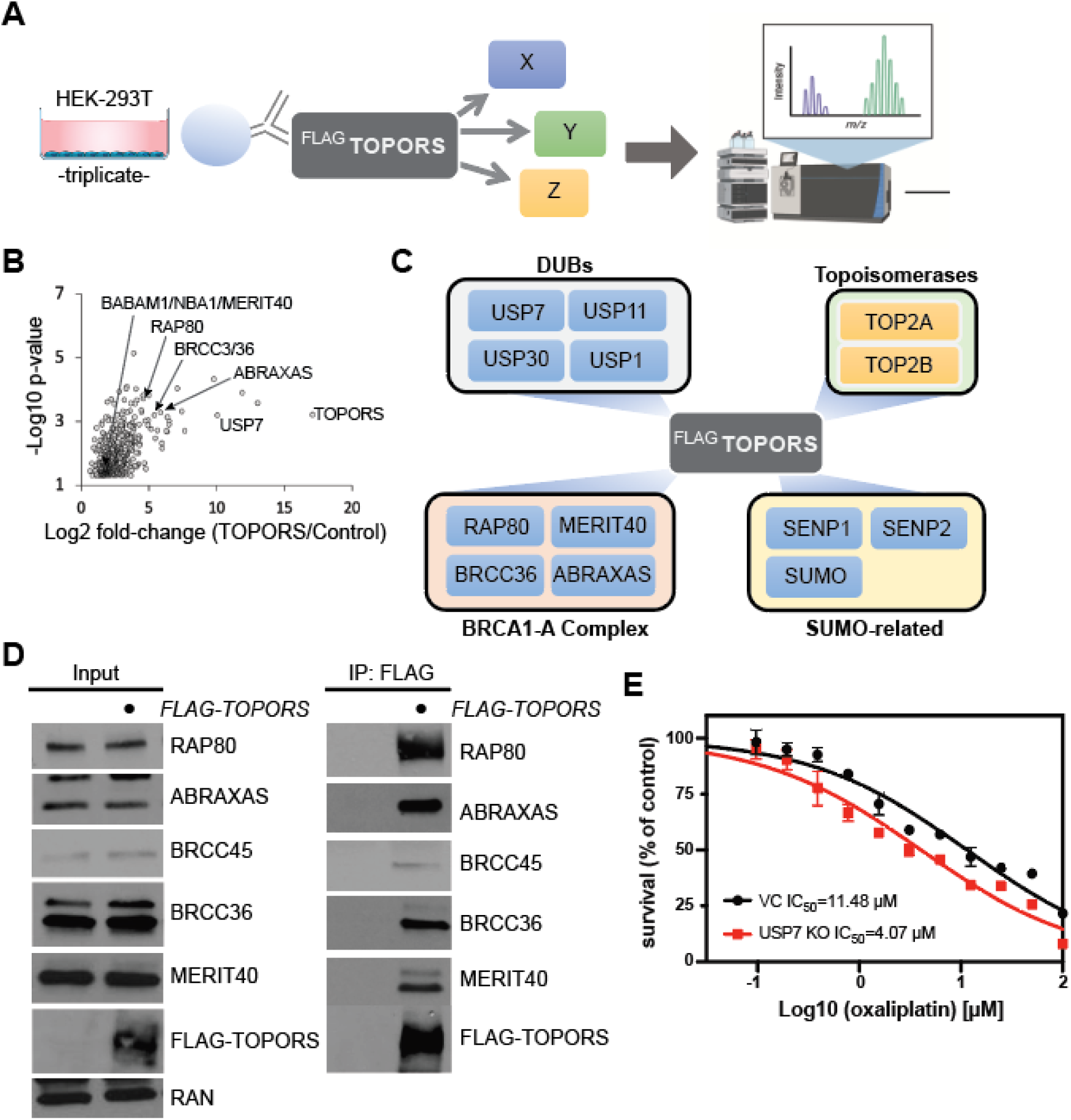
TOPORS interacts with the BRCA1-A complex. (A) TOPORS IP-MS schematic. FLAG-TOPORS was transfected into HEK293T cells in triplicate. FLAG-tagged TOPORS or an empty vector was immunoprecipitated and samples were analyzed by LC MS/MS. (B) Proteins enriched in TOPORS IPs compared to controls are shown. (C) TOPORS interactome showing selected interactor families identified by IP-MS. (D) HEK293T cells were transfected with FLAG-TOPORS or an empty vector control for 24 h, FLAG was immunoprecipitated and eluates analyzed by Western blot for the indicated proteins. (E) Dose response curve for MDA-MB-231 USP7 KO (red) or vector control (black) cells treated with oxaliplatin continuously for 72 h. Each value represents mean of 3 experiments ± SD.

Also, among the top interacting proteins were several components of the BRCA1-A complex (Figure 7C). BRCA1 is a central mediator of DNA repair through homologous recombination, and significantly, is mutated in hereditary breast cancers that often develop into TNBC (Toyama et al. 2008). Using the FLAG-TOPORS expressing HEK293T cells, we validated the interaction between TOPORS and all the know components of the BRCA1-A complex using immunoprecipitation followed by Western blot analysis (Figure 7D). We identified RAP80, BABAM1/NBA1/MERIT40, BRCC3/BRCC36, and ABRAXAS1 as top interacting proteins. We also identified BRE/BRCC45, another component of the complex, only in our TOPORS pulldown, although we were not able to accurately quantify its enrichment. Remarkably, while we identified all of the members of the BRCA1-A complex, we did not detect an interaction with BRCA1 itself, either by mass spectrometry or IP-western.

Collectively with published data, our results suggest a potential role for TOPORS in DNA damage repair. Interestingly, TOPORS depletion in SUM159 and MDA-MB-231 cells with siRNA had no significant impact on the levels of the BRCA1 complex proteins, suggesting that TOPORS might not control its degradation (Supplemental Figure 6A). Our data, and that of others, link TOPORS with critical cancer pathways, with evidence suggesting that its loss sensitizes cells to DNA-damaging agents (Hariharasudhan et al. 2022; Renner et al. 2010; Lin et al. 2005). To investigate whether USP7 inhibitors, which destabilize TOPORS, could enhance the efficacy of genotoxic chemotherapies, we tested the impact of USP7 inhibition in combination with DNA damaging drugs. While treatment with the USP7 inhibitor FT671 alone did not affect γH2A.X staining, co-treatment with FT671 and camptothecin reduced γH2A.X levels (Supplemental Figure 6B-C). Treatment of USP7 KO cells with oxaliplatin using the BRCA-proficient MDA-MB-231 cell lines significantly reduced the IC50 values compared to control cells. Specifically, the IC50 for oxaliplatin decreased from 11.48 µM in control cells to 4.07 µM in USP7 KO cells (Figure 7E). These findings suggest that USP7 inactivation sensitizes TNBC cells to DNA-damaging agents through TOPORS degradation and destabilization of the BRCA1-A complex.

## DISCUSSION

USP7, a deubiquitinase that stabilizes numerous pro-tumorigenic proteins, has emerged as a critical oncogene in cancer biology. Our findings, alongside prior research, establishes USP7 as a key driver of TNBC tumor growth and metastasis through its regulation of protein stability and downstream signaling pathways (Yi et al. 2023; Huang et al. 2024; Lin et al. 2022). Through genetic knockout and pharmacological inhibition, we demonstrated that targeting USP7 significantly impairs TNBC progression. Moreover, using deep quantitative proteomics we identified several novel USP7 regulated targets, including TOPORS. These findings expand our understanding of USP7’s role in TNBC progression and highlight its potential as a therapeutic target. Targeting USP7 could disrupt proteostasis and enhance the effectiveness of DNA-damaging therapies, offering a promising strategy for treating this aggressive breast cancer subtype.

In our broad analysis of breast cancer patient data, we found USP7 is notably overexpressed at the mRNA level and frequently amplified at the chromosomal level, making it one of the most recurrently upregulated DUBs in the disease. Its upregulation is observed in nearly one-quarter of breast tumors, surpassing even the prevalence of CMYC alterations. USP7 overexpression occurs across all major breast cancer subtypes, including ER-positive, HER2-positive, and TNBC tumors, and is independent of *TP53* mutation status. Thus, in line with recent reports, USP7 likely contributes to tumorigenesis through p53-independent mechanisms (Yi et al. 2023; Saha et al. 2022; Tavana et al. 2018). This broad expression pattern, coupled with the association between high USP7 levels and poor patient outcomes, highlights its importance in breast cancer pathogenesis and progression.

Our in vivo studies revealed that USP7 KO tumors exhibit significantly reduced growth and metastasis to lymph nodes, lung, and bones, leading to extended survival. These findings emphasize USP7’s role in driving tumor aggressiveness. Furthermore, the increased necrosis observed in USP7 KO tumors indicate heightened tumor stress or impaired adaptation to the tumor microenvironment, warranting further investigation into the interaction between USP7 and the tumor microenvironment in metastatic progression. Pharmacological inhibition of USP7 using the selective inhibitor FT671 produced effects comparable to genetic ablation, significantly impairing tumor growth without evident toxicity during treatment. These findings validate the clinical relevance of USP7 inhibitors as a therapeutic strategy in TNBC. Recently, additional strides have been made in the development of USP7 inhibitors, through various screening and chemical modification strategies (Futran et al. 2024; Leger et al. 2020; Vasas et al. 2024). Evaluation of these compounds ability to impede tumor growth, impact cancer phenotypes, and reshape proteomes in disease specific models, while considering potential toxicities, is a critical next step in evaluating the therapeutic potential of USP7 in TNBC.

Mechanistically, our multi-omics approach revealed USP7’s broad regulatory role in TNBC biology. Our proteomic analysis identified USP7 as a master regulator of TNBC proteostasis, controlling the stability of multiple E3 ligases, including TRIP12 (Cai et al. 2015), TRIM27 (Hao et al. 2015), and TOPORS. Notably, TOPORS emerged as a novel USP7 substrate with critical roles in cell proliferation, DNA damage response, and relieving DNA-protein crosslinks (Carnie et al. 2024; Liu et al. 2024; Rasheed et al. 2002; Hariharasudhan et al. 2022; Lin et al. 2005). While TOPORS’s function in TNBC has been unexplored, our findings suggest that USP7 stabilizes TOPORS, and its inhibition leads to TOPORS degradation. Given the association of TOPORS loss with increased sensitivity to DNA-damaging agents, our data point to a potential mechanism underlying the therapeutic synergy between USP7 inhibitors and genotoxic chemotherapy.

Collectively, these findings position USP7 as a central regulator of TNBC, influencing tumor biology, influencing critical pathways such as ubiquitination and DNA repair. The identification of TOPORS as a USP7 substrate expands our understanding of its functional network and provides additional therapeutic opportunities. The potential to combine USP7 inhibitors with DNA-damaging agents represents an exciting avenue for future research. Our results also suggest that protein-based markers could guide therapeutic selection in TNBC in the future, and this could include BRCA1-A complex members, TOPORS, or USP7 itself.

Further studies are necessary to fully elucidate the spectrum of USP7-regulated pathways and to explore its role across different TNBC subtypes. Understanding the molecular contexts and biomarkers that predict USP7 inhibitor efficacy will be crucial for translating these findings into clinical applications, ultimately improving outcomes for patients with TNBC. Future studies should aim to elucidate the full spectrum of USP7-regulated pathways and explore its role in other TNBC subtypes. Understanding the molecular context in which USP7 inhibition is most effective will be critical for translating these findings into clinical applications.

## MATERIALS AND METHODS

### Selection of TNBC cases from METABRIC and survival data

Survival annotated data from 2509 patients with breast cancer was obtained from the METABRIC (Curtis et al. 2012) study using cBioPortal on May 17, 2023. We identified 199 patients with TNBC with associated mRNA microarray data. The impact of USP7 expression on overall survival was assessed by stratifying patients into quartiles (Q1-Q4) based on USP7 z-score expression level. Kaplan-Meier curves were plotted using the R package survival and statistical significance was assessed using the log-rank test and defined as p<0.05.

Overall survival information was provided in the data_clinical_patient.txt file for all 199 patients IDs under OS_STATUS and VITAL_STATUS. We analyzed 10-year survival (120 months) with “Died of disease” cases flagged as events while “Living” and “Died of Other Causes” cases were censored observations. The gene expression data for these 199 patients was already associated with HUGO gene symbols by cBioPortal. Samples were separated into four quartiles, Q1 thru Q4 based on USP7 z-score expression.

#### Immunohistochemistry

Tissue microarray (TMA) slides containing 110 TNBC cases were obtained from US Biomax (BRE1201; TissueArray.com). Slides were baked for 2 h at 60°C, de-paraffinized with xylene, washed with ethanol, and hydrated. Slides were sub-boiled using 1X sodium citrate for 10 min and cooled. Sections were incubated with 3% hydrogen peroxide for 15 min, washed, and blocked using 5% goat serum in TBS-0.25% Tween (TBS-T) for 1 h at room temperature. USP7 (4833S; Cell Signaling Technology) primary antibody dilution (1:100 2% goat serum in TBS-T) was incubated overnight at 4°C. Next day, slides were washed three times with TBS-T and incubated with 1:200 dilution of goat anti-rabbit biotinylated in 2% goat serum in TBS-T for 30 min at room temperature. Slides were washed three times with TBS-T and incubated for 30 min at room temperature with avidin/biotin-based peroxidase system (PK-6100; Vector Laboratories). After slides were washed three times with TBS-T, 3,3-diaminobenzidine (DAB) was added to stain slides for 2 min, followed by 10 sec of hematoxylin counterstain. Slides were mounted using Gelvatol and scanned using Ventana DP200. All histological analysis was conducted using QuPath.

#### Cell culture

TNBC cell lines MDA-MB-231, MDA-MB-468, and SUM159, as well as the HER2-negative breast cancer cell line MCF7 and the immortalized normal breast cell line MCF10A, were acquired from the American Type Culture Collection (ATCC). TNBC cell lines and MCF7 cells were cultured in Dulbecco’s modified Eagle’s medium (DMEM) high glucose (Gibco) and 10% fetal bovine serum (FBS; Gibco) with penicillin/streptomycin (P/S; Gibco). MCF10A cells were cultured in DMEM high glucose + 10% FBS with P/S and 20 μg/ml epidermal growth factor (EGF; Sigma-Aldrich). HEK293T cells were purchased from ATCC and cultured in high-glucose DMEM supplemented with 10% FBS and P/S. All cell lines were maintained in a humidified incubator at 37°C with 5% CO_2_ and regularly screened for mycoplasma using a mycoplasma detection kit (13100-01, SouthernBiotech). All cell lines were routinely confirmed by STR profiling at the Arizona Genetics Core.

#### Western blotting

Cell pellets or tumor tissues were lysed for 30 min on ice for protein extraction using RIPA buffer (50 mM Tris-HCl (pH 8), 150 mM NaCl, 1% NP-40, 0.5% sodium deoxycholate, 0.1% SDS, 1mM EDTA) with protease inhibitor (100X Halt Protease Inhibitor Cocktail; Thermo Fisher). Lysates were centrifuged at 14000 rpm for 30 min at 4°C. The total protein concentration in the supernatant was quantified using a BCA protein assay (Pierce BCA Protein Assay Kit). The same concentration of total protein for lysates was loaded and resolved on a 4-15% gradient SDS-PAGE gel (Bio-Rad). Proteins were transferred onto a PVDF membrane (EMD Millipore) by semi-dry transfer. The membrane was incubated for 1 h at room temperature with 3% non-fat dry milk in TBS-T to block non-specific binding. Primary antibody was diluted in 0.5% non-fat dry milk in TBS-T at: 1:1000 USP7 (4833S; Cell Signaling Technology) and 1:4000 β-actin (A2228; Sigma) and incubated overnight at 4°C on a shaker. Following incubation, the membrane was washed twice with TBS-T and incubated with either goat anti-rabbit (PI-1000-1; Vector Laboratories) or horse anti-mouse (PI-2000-1; Vector Laboratories) HRP-conjugated secondary antibody diluted at 1:5000 in 0.5% non-fat dry milk in TBS-T and incubated for 30 min with shaking at room temperature. Membrane was washed twice with TBS-0.25% Tween, and bands were visualized using a chemiluminescent substrate (34580; Thermo Scientific) and imaged using Syngene G:BOX. The bands were quantified using ImageJ software.

Alternatively, cells were harvested in 1x PBS by gently scraping. The cells were pelleted by centrifuging at 3500 rpm for 5 mins. The cell pellets were lysed in NETN (20 mM Tris pH 8.0, 100 mM NaCl, 0.5 mM EDTA, 0.5% NP40) lysis buffer freshly supplemented with protease (1mg/mL aprotinin, 1mg/mL leupeptin, 1mg/mL pepstatin) and phosphatase (25mM Na3VO4, and 25mM ABESF) inhibitors for 15 min on ice. The lysate was clarified by centrifuging at 15000 rpm for 15 min. The total protein in the clarified cell lysate was measured using Bradford assay (Bio-rad). 25 μg of protein was separated using 4-20% gradient gel (Bio-rad) and proteins electroblotted to nitrocellulose membrane at 100V for 1 h. The blot was blocked for an hour, rinsed, and probed with appropriate antibodies overnight at 4°C. The blots were then rinsed in TBS-T three times followed by incubation with an HRP-conjugated secondary antibody against respective host for an hour at room temperature. The blots were washed with TBS-T three times, and the blot was incubated in ECL for 2 mins. The chemiluminescence was captured using Amersham Hypefilms and developed. The primary antibodies used were: 1:1000 TOPORS (A302-179A; Bethyl), 1:1000 USP7 (sc-30164; Santa Cruz), 1:1000 Rap80 (14466S; Cell Signaling Technology), 1:500 Abraxas (A302-180A; Bethyl), 1:1000 Merit40 (12711; Cell Signaling Technology), 1:1000 BRCC36 (18215; Cell Signaling Technology), BRCC45/BRE (12457; Cell Signaling Technology). 1:1000 Tubulin (sc-32293; Santa Cruz) or 1:1000 Ran (sc-271376; Santa Cruz) was used as loading control.

#### Immunoprecipitation

HEK293T cells were transected with 2μg of plasmid encoding either Flag-TOPORS or Myc-USP7 or both using PolyJet (SignaGen). The transfection was performed overnight, followed by a media change and the cells were incubated for another 24 hours. The total length of transfection was 48hrs. The cells were harvested on ice by scraping in 1x PBS. The harvested cells were pelleted and lysed in NETN lysis buffer. A protein quantification using Bradford assay (Bio-rad) was performed. To immunoprecipitate TOPORS or USP7, Flag or Myc beads was used respectively. The beads (Sigma) were prepared by washing with PBS-T twice and then with cold NETN buffer. Equal amounts of protein from all groups were used for immunoprecipitation. The protein lysate and beads were nutated at 4°C for an hour. The beads were collected by centrifugation, washed with cold lysis buffer three times. After the final wash, the bound protein was eluted using 2X Laemelli buffer by boiling at 100 °C for 5 mins. The sample was then separated using SDS-PAGE.

#### siRNA transfection

siRNA against TOPORS (siTOPORS sequence 1: GGUCUCGUAGCAGUGAUCAUGGUAA; siTOPORS sequence 2: UUACCAUGAUCACUGCUACGAGACC; Invitrogen) or USP7 (siUSP7 #1: CUAAGGACCCUGCAAAUUA; siUSP7 #2: GUGGUUACGUUAUCAAAUA; siUSP7 #3: UGACGUGUCUCUUGAUAAA; siUSP7 #4: GAAGGUACUUUAAGAGAUC; Sigma) was transfected using Lipofectamine RNAiMAX (Invitrogen) using the manufacturer’s protocol. Briefly, the cells were plated overnight before the transfection at a lower density. The siRNA and RNAiMAX was diluted to appropriate concentrations in OptiMEM (Invitrogen) in two separate tubes. The diluted RNAiMAX was added to the siRNA tube and incubated at room temperature for 15mins before adding to the cells. Before making the transfection mix, the culture media in the cells were changed to basal DMEM. The cells with the transection mix allowed to grow overnight. The next day, media with the transfection mix was replaced with complete DMEM culture media and cells were allowed to grow for another 24 hours. An immunoblot was performed to confirm the depletion of either TOPORS or USP7.

#### Lentivirus production and transduction

*USP7* gRNA (gRNA1 sequence: AGATGTATGATCCCAAAACG and gRNA2 sequence: GTGTACATGATGCCAACCGA) was cloned into the doxycycline-inducible CRISPR/Cas9 plasmid TLCV2 (87360; Addgene). pLenti PGK V5-LUC Neo (21471; Addgene) was used for bioluminescence imaging. All lentiviruses were produced by calcium phosphate transfection using HEK293T cells and the lentiviral TLCV2 transfer vector, lentiviral envelope plasmid pCMV-VSV-G (8454; Addgene), and lentiviral packaging plasmid pCMV-dr8.2-dvpr (8455; Addgene). Supernatants containing lentiviral particles were collected 24 and 48 h post-transfection and centrifuged at 1600 g for 10 min at 4°C to remove cell debris. Supernatants were filtered through 0.45 μm PES; filter (ge-0504; Nalgene) and concentrated by ultracentrifugation at 23000 rpm for 1.5 hours at 4°C. Lentiviral pellets were resuspended in cold PBS and stored in aliquots at −80°C. Following lentiviral titering, transduction of target cells was achieved by exposing cells to viral particles in serum-free condition for 6 h using a 0.2 MOI. Selection was carried using either puromycin or G418, depending on the vector.

#### CRISPR KO single cloning

The CRISPR KO pool of clones was generated by inducing SUM159, MDA-MB-231, and MDA-MB-468 TLCV2 *USP7* gRNA cells with 1 μg/ml doxycycline. 24- and 48-hours post-induction, cells were checked for GFP expression, plated at a low density of 100-200 cells per 100 mm dish, and were allowed to expand as single colonies for 3 weeks. The cell medium was removed from the dish, and 2% agarose with 2X PBS mixed in equal parts was poured onto the dish to solidify. Trimmed p200 pipette tips were used to puncture and draw up agarose and were vigorously pipetted up and down into a low volume of cell medium in each well, generating bubbles. The dispersed cells were allowed to adhere, recover, and expand as single clones. After numerous rounds of cell proliferation, single clones were screened for KO status using Western blotting. Single clones with the lowest protein expression levels of USP7 were selected for in vitro and in vivo studies. KO clones were assessed for CRISPR genome editing using Tracking of Indels by Decomposition (TIDE).

#### Cell proliferation, colony formation assay

Cells were seeded at 10-20% confluency in 96-well plates and imaged every 24 h using a BioTek BioSpa and Cytation5 cell imaging reader. Cell proliferation was assessed by cell count quantification using ImageJ software.

For the soft-agar colony formation assay, 3000 cells were mixed in 0.36% agarose with full serum media, and the top layer was allowed to solidify above the bottom layer containing 0.75% agarose in full media. Single cells were given five weeks to form colonies. Colonies embedded in agarose were fixed and stained with 0.1% crystal violet in 10% ethanol at room temperature for 15 min. Agarose was de-stained in multiple rounds of washing with Milli-Q water and imaged using a BZ-X700 Keyence microscope. The colonies were manually counted under a light microscope.

USP7 inhibitors, XL177A or FT671, were used at 1 μM (for SUM159 and MDA-MB-231) or 5 μM (for MDA-MB-468) and replenished every 24 hours.

#### Chemicals

USP7 inhibitors: XL177A (10 mM in DMSO) was kindly provided by Sara J. Buhrlage (Schauer et al. 2020). Stock solution was aliquoted and stored at −20°C. FT671 was purchased from MedChemExpress (cat# HY-107985). Stock solution of 10 mM in DMSO was prepared, aliquoted, and stored at −80°C.

#### Scratch-wound healing assay

Cells were seeded as a monolayer in 24-well plates the day before the scratch to reach 70-80% confluency. 2 h prior scratch, cells were treated with 5 μg/ml mitomycin C (S8146; Sigma) to control for proliferation differences. Using a 200 μl pipette tip, the monolayer was scratched across the center of the well. Images were captured immediately (0 h) and 8 h after the wound at the same location. The distance between the boundaries of the wound at 0 and 8 h at 10 different locations were measured using ImageJ software. Assay was performed in triplicate, with each bar representing mean ± standard deviation. For the migration assays with USP7 inhibitors, 1 μM (for SUM159 and MDA-MB-231) or 5 μM (for MDA-MB-468) XL177A or FT671 was added to the cells immediately after the scratch (0 h).

#### Transwell invasion assay

Cells were treated with 5 μg/ml mitomycin C for 2 h at 37°C. 8 μm pore PET membranes (0877121; Thermo Fisher) were coated with 25 μg of Matrigel per insert and were incubated for 1 h at 37°C to solidify prior to adding a layer of cells resuspended in serum-free media. Full-serum medium containing 100 ng/ml FGF-2 was added to the wells beneath the inserts. Cells were allowed to invade the Matrigel overnight at 37°C to the bottom side of the insert. The remaining cells in the inner chamber were swabbed, and cells on the bottom side of the insert were fixed with 70% ethanol and stained with 0.2% crystal violet. The inserts were de-stained with Milli-Q water and imaged using BioTek Cytation5. The images were stitched using the ImageJ software, and the stained cells were manually counted. Assay was performed in triplicates, with each bar representing mean ± standard deviation. For the invasion assays with USP7 inhibitors, 1 μM (for SUM159 and MDA-MB-231) or 5 μM (for MDA-MB-468) XL177A or FT671 was added with the cells onto the Matrigel.

#### Animal studies

NOD *scid* gamma (NSG) mice were acquired from Jackson Laboratory. 8-to-10-week-old NSG female mice were injected orthotopically under the mammary fat pad located below the fourth nipple with 50 μl of 2 × 10^6^ cells resuspended in sterile 1X PBS. Surgery was performed on anesthetized and shaved mice. Using tweezers to lift the skin around the nipple, the fat pad was visible under light. A 27G tuberculin syringe with an attached needle was then used to enter the skin a few millimeters from the center of the nipple, and cells were slowly injected under the center. After the needle was slowly removed, the injection site was observed for signs of leakage. The mice were removed from anesthesia and allowed time to recover post-surgery. Tumor sizes were measured biweekly either by caliper measurement or bioluminescence imaging (IVIS Lumina). Primary tumors, lungs, and visible metastatic tumors were collected and used for analysis. To confirm the presence of metastasis, ex vivo imaging was conducted on the lymph nodes, lungs, liver, and bones.

For oral gavage, FT671 was dissolved in 10% DMSO/90% PEG400. Mice received a lead-in dose of 25 mg/kg of FT671 or vehicle when the tumors had reached an average volume of 168 mm^3^. Mice were then dosed daily for 20 days at 50 mg/kg of FT671 or vehicle. At the study endpoint, primary tumors and lungs were collected for analysis. Blood was collected by cardiac puncture; 10 μl was immediately mixed in with 10 μl 100 mM EDTA and analyzed for complete blood count, and the remaining blood was left to clot for 30 min at room temperature. The blood sample was spun at 2000 x g for 10 min at 4°C to collect the supernatant (serum). Serum was analyzed via Standard Tox Panel (62794; IDEXX BioAnalytics).

The University of California Irvine Institutional Animal Care and Use Committee approved all animal procedures.

#### RNA sequencing

Total RNA was isolated using RNA Microprep Kit (R1050; Zymo Research) or RNeasy Mini Kit (74104; Qiagen). 500 ng of total RNA was used to enrich for poly(A) mRNA to generate RNA sequencing library using NEBNext Ultra II Directional RNA Library Kit for Illumina per manufacturer’s protocol (E7760L; New England Biolabs). Samples were purified with AMPure XP beads (A63881; Beckman Coulter) and eluted in 20 μl of 0.1X TE buffer. Samples were checked for library quality using Agilent BioAnalyzer 2100 and quantified using KAPA qPCR. A paired-end 100-bp sequencing run was conducted on Illumina NovaSeq X Plus with 50 million PE reads.

#### Proteomics

SUM159, SUM149, and MDA-MB-231 cells were treated with either DMSO, 1 μM FT671, or 1 μM XL177A for 8 hours prior to harvesting for global quantitative proteomics at the UNC Proteomics Core Facility. Briefly, treated cells were harvested and washed with ice cold 1X PBS three times. Cells were kept on ice or chilled throughout. Next, cells were lysed using urea lysis buffer (8 M Urea, 75 mM NaCl, 50 mM Tris, pH 8.0, 1 mM EDTA, 1X protease inhibitor cocktail and 1X phosphatase inhibitor cocktail) for 10 min on ice and were subsequently snap frozen in liquid nitrogen. Cell lysates were spun down the remove debris for 15 min at maximum speed in a refrigerated microcentrifuge. Protein concentration was determined by Bradford Assay. Lysates were stored at −80°C prior to submitting for analysis to the Proteomics Core. All analyses were performed in triplicates.

TMT Proteomics Analysis. Sample Preparation: Protein lysates (300 µg) were reduced with 5mM DTT for 45 min at 37°C and alkylated with 15mM iodoacetamide for 30 min in the dark at room temperature. Samples were digested with LysC (Wako, 1:50 w/w) for 2 hr at 37°C, then diluted to 1M urea and digested with trypsin (Promega, 1:50 w/w) overnight at 37°C. The resulting peptide samples were acidified to 0.5% trifluoracetic acid, desalted using desalting spin columns (Thermo), and the eluates were dried via vacuum centrifugation. Peptide concentration was determined using a Fluorometric Peptide Assay (Pierce). Samples were split into three sets (n = 9 each) based on cell line, and labeled with TMT10plex reagents (Thermo Fisher). A pooled sample, consisting of ∼2µg of each sample were used to fill the extra TMT channel. 40 µg of each sample was reconstituted with 50 mM HEPES pH 8.5, then individually labeled with 150 µg of TMT reagent for 1 hr at room temperature. Prior to quenching, the labeling efficiency was evaluated by LC-MS/MS analysis of the pooled sample. After confirming >98% efficiency, samples were quenched with 50% hydroxylamine to a final concentration of 0.4%. Labeled peptide samples were combined 1:1, desalted using Thermo desalting spin column, and dried via vacuum centrifugation. The two dried TMT-labeled samples were fractionated using high pH reversed phase HPLC. Briefly, the samples were offline fractionated over a 90 min run, into 96 fractions by high pH reverse-phase HPLC (Agilent 1260) using an Agilent Zorbax 300 Extend-C18 column (3.5-µm, 4.6 × 250 mm) with mobile phase A containing 4.5 mM ammonium formate (pH 10) in 2% (vol/vol) LC-MS grade acetonitrile, and mobile phase B containing 4.5 mM ammonium formate (pH 10) in 90% (vol/vol) LC-MS grade acetonitrile. The 96 resulting fractions were then concatenated in a non-continuous manner into 24 fractions and dried down via vacuum centrifugation and stored at −80°C until further analysis.

#### LC/MS/MS Analysis

The samples consisting of 2 sets of 24 fractions (48 fractions total) were analyzed by LC/MS/MS using an Ultimate 3000-Exploris480 (Thermo Scientific). Samples were injected onto an IonOpticks Aurora series 2 C18 column (75 μm id × 15 cm, 1.6 μm particle size), and the samples were separated over a 140 min method. The gradient for separation consisted of 5–45% mobile phase B at a 250 nl/min flow rate, where mobile phase A was 0.1% formic acid in water and mobile phase B consisted of 0.1% formic acid in 80% acetonitrile. The Exploris480 was operated in turboTMT10 mode with a cycle time of 3s. Resolution for the precursor scan (m/z 375–1400) was set to 60,000 with an AGC target set to 100%; maximum injection time set to auto. Resolution of the MS2 scan was set to 30,000, and consisted of HCD normalized collision energy (NCE) 33; AGC target set to 300%; maximum injection time of 55 ms; isolation window of 0.7 Da.

#### TMT Proteomics Data Analysis

Raw data were processed using Proteome Discoverer (version 2.5) for peptide/protein identification and TMT10plex quantitation. Data were searched against a Uniprot Reviewed Human database (downloaded 01/2022, containing 20,360 sequences), appended with a common contaminants database, using the Sequest HT search engine. A maximum of two missed tryptic cleavages were allowed, and fixed modifications were set to TMTpro peptide N-terminus and Lys and carbamidomethyl Cys. Dynamic modifications were set to N-terminal protein acetyl and oxidation Met. Precursor mass tolerance was set to 10 ppm, and fragment mass tolerance was set to 0.02 Da. MS2 coisolation interference threshold was set to 30%. The peptide false discovery rate was set to 1%. Reporter ion abundance was calculated based on intensity. Results were filtered to 5% protein FDR, a minimum of 2 peptides, and quantitation in minimum 50% of samples. Statistical analysis (Student’s T-test, unpaired) was conducted within Argonaut (Mertins et al. 2018). Proteins with an absolute log2 fold change ≥ 0.5 and a p-value < 0.05 are considered significant. The mass spectrometry proteomics data have been deposited to the ProteomeXchange Consortium via the PRIDE partner repository (Perez-Riverol et al. 2024) with the dataset identifier PXD059902.

### Reviewer access details

Log in to the PRIDE website using the following details:

**Project accession:** PXD059902

**Token:** e1bjASE13is4

Alternatively, reviewer can access the dataset by logging in to the PRIDE website using the following account details:

**Password:** Aa5vwSRtW45L

#### Sample Preparation for Affinity Purification Mass Spectrometry Analysis (AP-MS)

Immunoprecipitated samples from FLAG-TOPORS or empty vector control pulldowns, performed in triplicate in HEK293T cells (one 10cm dish per sample), were subjected to on-bead trypsin digestion, as previously described (Mouery et al. 2024). After the last wash buffer step, 50 µl of 50 mM ammonium bicarbonate (pH 8) containing 1 µg trypsin (Promega) was added to beads overnight at 37°C with shaking. The next day, 500 ng of trypsin was added then incubated for an additional 3 h at 37°C with shaking. Supernatants from pelleted beads were transferred, then beads were washed twice with 50 µl LC/MS grade water. These rinses were combined with original supernatant, then acidified to 2% formic acid. Peptides were desalted with peptide desalting spin columns (Thermo) and dried via vacuum centrifugation. Peptide samples were stored at −80°C until further analysis.

#### LC/MS/MS Analysis for TOPORS AP-MS

Each sample was analyzed by LC-MS/MS using an Easy nLC 1200 coupled to a QExactive HF (Thermo Scientific). Samples were injected onto an IonOpticks Aurora Elite TS C18 column (75 μm id × 15 cm, 1.7 μm particle size) and separated over a 120 min method. The gradient for separation consisted of a step gradient from 5 to 36 to 48% mobile phase B at a 250 nl/min flow rate, where mobile phase A was 0.1% formic acid in water and mobile phase B consisted of 0.1% formic acid in 80% ACN. The QExactive HF was operated in data-dependent mode where the 15 most intense precursors were selected for subsequent HCD fragmentation. Resolution for the precursor scan (m/z 350–1700) was set to 60,000 with a target value of 3 × 106 ions, 100ms inject time. MS/MS scans resolution was set to 15,000 with a target value of 1 × 105 ions, 75ms inject time. The normalized collision energy was set to 27% for HCD, with an isolation window of 1.6 m/z. Peptide match was set to preferred, and precursors with unknown charge or a charge state of 1 and ≥ 8 were excluded.

#### Data Analysis for TOPORS AP-MS

Raw data were processed using the MaxQuant software suite (version 1.6.15.0) for peptide/protein identification and label-free quantitation (Tyanova et al. 2016a). Data were searched against a Uniprot Reviewed Human database (downloaded 01/2023, containing 20,404 sequences), appended with the HepaCAM sequences, using the integrated Andromeda search engine. A maximum of two missed tryptic cleavages were allowed. The variable modifications specified were: N-terminal acetylation and oxidation of Met. Label-free quantitation (LFQ) was enabled. Results were filtered to 1% FDR at the unique peptide level and grouped into proteins within MaxQuant. Match between runs was enabled. Data filtering and statistical analysis was performed in Perseus software (version 1.6.14.0) (Tyanova et al. 2016b). The mass spectrometry proteomics data have been deposited to the ProteomeXchange Consortium via the PRIDE partner (Perez-Riverol et al. 2024) repository with the dataset identifier PXD059910.

### Reviewer access details

Log in to the PRIDE website using the following details:

**Project accession:** PXD059910

**Token:** etZGC48UXui6

Alternatively, reviewer can access the dataset by logging in to the PRIDE website using the following account details:

**Password:** 4qoneFDhZGoT

#### Statistical analysis

All graphs and statistical analyses were performed using the GraphPad Prism software (Prism 10 for macOS, GraphPad Software). The sample size (*n*), statistical tests, and *p-*values are noted in the figure legends. Results are presented as the mean ± SD. Statistical significance of differences between two groups in vitro was assessed by Student’s *t*-test, with *p*-values < 0.05 statistically significant. A two-way ANOVA was conducted for animal studies. Blind analysis was performed for the phenotypic measurements.

## ACKNOWLEDGMENTS

We thank lab members and colleagues at UNC and UCI for helpful discussions throughout this.

The Emanuele lab is supported by the UNC University Cancer Research Fund, NIGMS R35GM153250, ACS RSG-18-220-01-TBG and R01CA280482. TOPORS expression clones were kindly provided by Lienhard Schmitz (Justus-Liebig-University). The Benavente lab is supported by NCI (R01CA229696), American Cancer Society (RSG-19-031-01-DMC, ACS-05609104), and the Chao Family Comprehensive Cancer Center A1 Bridge Funding. The Spanheimer lab is supported by K08CA280388, R37CA292075, and R01CA280482. The UNC Proteomics Core Facility is supported in part by NCI Center Core Support Grant (2P30CA016086-45) to the UNC Lineberger Comprehensive Cancer Center. M.S-P. is supported by the Fulbright Scholar Program, which is sponsored by the U.S. Department of State and the Comisión Fulbright España. This work was also made possible, in part, through access to the Genomics High Throughput Facility Shared Resource of the Cancer Center Support Grant (P30CA-062203) at the University of California, Irvine and NIH shared instrumentation grants 1S10RR025496-01, 1S10OD010794-01, and 1S10OD021718-01.

## AUTHOR CONTRIBUTIONS

M.J.E. and C.A.B. conceived the project. A.K. performed most cell-based and in vivo experiments. P.G. performed most of the in vitro assays. N.U. and M.S-P. assisted with in vitro and cell-based experiments. A.M., N.K.B., L.E.H. assisted in designing proteomics experiments and performed and helped analyze data for all mass spectrometry experiments. C.C.C. performed RNA-seq data analysis. R.T.K. and P.M.S. analyzed data from METABRIC and TCGA databases. A.K., M.J.E., and C.A.B. discussed and wrote the manuscript. All authors provided feedback on the final draft.

